# Asymmetrical interference between number and item size perception provide evidence for a domain specific impairment in dyscalculia

**DOI:** 10.1101/332155

**Authors:** Elisa Castaldi, Anne Mirassou, Stanislas Dehaene, Manuela Piazza, Evelyn Eger

**Affiliations:** Cognitive Neuroimaging Unit, CEA DRF/I2BM, INSERM, Université Paris-Sud, Université Paris-Saclay, NeuroSpin center, France; Centre Hospitalier Rives de Seine, Service de Pédiatrie et Néonatologie, Unité de Dépistage des troubles des apprentissages, France; Center for Mind/Brain Sciences, University of Trento, Italy

**Keywords:** Numerical cognition, Numerosity perception, Mean size perception, Developmental dyscalculia, Inhibitory control

## Abstract

Dyscalculia, a specific learning disability that impacts arithmetical skills, has previously been associated to a deficit in the precision of the system that estimates the approximate number of objects in visual scenes (the so called ‘number sense’ system). However, because in tasks involving numerosity comparisons dyscalculics’ judgements appears disproportionally affected by continuous quantitative dimensions (such as the size of the items), an alternative view linked dyscalculia to a domain-general difficulty in inhibiting task-irrelevant responses.

To arbitrate between these views, we evaluated the degree of reciprocal interference between numerical and non-numerical quantitative dimensions in adult dyscalculics and matched controls. We used a novel stimulus set orthogonally varying in mean item size and numerosity, putting particular attention into matching both features’ perceptual discriminability. Participants compared those stimuli based on each of the two dimensions. While control subjects showed no significant size interference when judging numerosity, dyscalculics’ numerosity judgments were strongly biased by the unattended size dimension. Importantly however, both groups showed the same degree of interference from number when judging mean size. Moreover, only the ability to discard the irrelevant size information when comparing numerosity (but not the reverse) significantly predicted calculation ability across subjects.

Overall, our results show that numerosity discrimination is less prone to interference than discrimination of another quantitative feature (mean item size) when the perceptual discriminability of these features is matched, as here in control subjects. By quantifying, for the first time, dyscalculic subjects’ degree of interference on another orthogonal dimension of the same stimuli, we are able to exclude a domain-general inhibition deficit as explanation for their poor / biased numerical judgement. We suggest that enhanced reliance on non-numerical cues during numerosity discrimination can represent a strategy to cope with a less precise number sense.

## Introduction

Evaluating how many objects are in a visual image requires disambiguating the discrete number of items from different continuous quantities, such as total contrast and luminance, area, density, and so on. A longstanding and influential theory in the field of numerical cognition proposes that humans are born with a ‘number sense’ (Dehaene, 1997; for a review see: Nieder, 2016), a phylogenetically ancient ability to make spontaneous and rapid estimates of the approximate number of objects in a visual scene. However, if covarying continuous features already provide cues from which numerosity can be inferred, behavioral performance might not be based on a specific sense of number. Previous research has addressed this issue by making non-numerical cues uninformative for numerosity decisions and successfully demonstrated that numbers can still be perceived, even from very early on in life (Brannon et al., 2004; Cordes and Brannon, 2011, 2011; de Hevia et al., 2017; Libertus et al., 2014; Piazza et al., 2004; Xu, 2003; Xu and Spelke, 2000). At the neuronal level, the brain structures found to be most involved in numerosity representation also seem to code for number independently of other perceptual dimensions. Indeed both neuroimaging experiments in human adults and children as well as monkey neurophysiology showed evidence for number-related neural signatures with a considerable level of generalization across other quantities and independence from low-level factors of the image (Cantlon et al., 2006; Castaldi et al., 2016; Eger et al., 2009; Fornaciai et al., 2017; Harvey et al., 2013; Harvey and Dumoulin, 2017; Izard et al., 2008; Nieder, A. et al., 2002; Nieder and Merten, 2007; Nieder and Miller, 2004; Piazza et al., 2004).

Despite much behavioral, neurophysiological and neuroimaging evidence suggesting that numerosity can be perceived directly through dedicated neuronal mechanisms (for reviews on the respective fields see: Anobile et al., 2016; Nieder, 2016; Piazza and Eger, 2016), both adults’ and children’s behavioral performance in numerosity tasks is often strongly affected by different combinations of covarying non-numerical quantities when these provide information of a direction incongruent with numerosity (Dakin et al., 2011; DeWind et al., 2015; Gebuis et al., 2009; Gebuis and Reynvoet, 2012a, 2012b; Hurewitz et al., 2006; Nys and Content, 2012; Ross, 2003; Rousselle et al., 2004; Rousselle and Noël, 2008; Salti et al., 2016; Sophian and Chu, 2008; Dénes Szűcs et al., 2013; Tokita and Ishiguchi, 2010). The underlying causes of this behavioral interference are not entirely understood, and several potential explanatory mechanisms have been proposed. One theory, prevailing in experimental psychology, is that different features of the stimulus are independently and automatically extracted, and compete for control of behavior (as in the classical STROOP effect, see for example Barth, 2008; Hurewitz et al., 2006; Nys and Content, 2012; Rousselle and Noël, 2008). This theory places the origin of interference at the level of the response selection. Alternatively, it has been proposed that interference may originate at the level of sensory extraction: models based on the stimulus energy at different spatial scales can yield non-veridical estimates of the number of items in a display resembling the biases of human observers (Dakin et al., 2011), and within hierarchical generative networks, interference from non-numerical quantities has been related to the efficiency of a normalization process embedded into the extraction of numerosity representations (Cappelletti et al., 2014b; Stoianov and Zorzi, 2017). Nevertheless, some authors have interpreted interference to indicate that numerosity is indirectly inferred from a combination of non-numerical quantitative features (though without specifying which combination of features in detail), sometimes going as far as to completely deny the existence of a dedicated perceptual mechanisms for numerosity (for a review see: Leibovich et al., 2016a).

It is noteworthy that among the studies that found strong interference of non-numerical dimensions on numerosity comparison, many required participants to judge rather difficult numerical ratios, even between 0.9 and 1.1 (DeWind et al., 2015; Nys and Content, 2012; Tokita and Ishiguchi, 2010). Importantly, the strongest interference is usually observed for the most difficult numerical ratios with a tendency to decrease for the easier comparisons (Hurewitz et al., 2006; Nys and Content, 2012). It is well-known that comparative judgments without counting are not perfect but approximate, depending on the ratio of the compared numbers with a precision that is commonly operationalized by the Weber fraction. It is hence conceivable that when subjects are required to make decisions close to or beyond the precision of their numerosity processing system, they would attempt to rely on associated quantities to solve the task, especially since in everyday life these often provide correlated information. However, such heuristic use of non-numerical information need not be the only possibility: even in symbolic number-size interference tasks, which are not limited by sensory/perceptual precision to the same extent as non-symbolic numerosity, the relative ratios of difference in the two dimensions predicted whether size interfered with number (Algom et al., 1996).

Despite the important role of relative discriminability and salience of the attended and unattended dimensions in interference paradigms, studies reporting interference from continuous dimensions onto non-symbolic numerical judgments have often neglected this aspect and paired difficult numerical ratios to be compared with often much larger differences in non-numerical quantities (e.g. Gebuis and Reynvoet, 2012a, 2012b, 2011; Hurewitz et al., 2006; Tokita and Ishiguchi, 2010). In sum, both the difficulty of the numerical ratio tested as well as the saliency of the unattended dimension with respect to the attended one may have contributed to the variations in the strength of interference described in the literature.

Compared to the wealth of studies on interference from other quantities on numerosity comparison, relatively fewer studies have investigated interference of numerosity onto judgement of a non-numerical quantitative dimension, most often total surface area (Barth, 2008; Hurewitz et al., 2006; Leibovich et al., 2016b; Nys and Content, 2012; Salti et al., 2016). These studies have to some extent arrived at different conclusions, sometimes finding that numerosity, and sometimes that area judgement is more subject to interference, possibly as a consequence of the above mentioned factor of degree of change / discriminability. Indeed when total surface area was claimed to be dominant over the numerical dimension, larger changes in the unattended area dimension were used (Hurewitz et al., 2006; Leibovich et al., 2016b), however when the range of ratio variation across dimension was physically equated, the opposite conclusion was reached (Nys and Content, 2012; Salti et al., 2016). Indeed the interference arising from numerosity changes in total surface area comparisons was reported to be either similar or stronger with respect to the total surface area interference during numerosity judgments, both when testing the subitizing range (Salti et al., 2016) and much higher numerosities (Nys and Content, 2012). However, none of these studies took into account the differences that may exist between the perceptual discriminability of different features, as a result of which using identical physical ratios across dimensions may not necessarily translate into equating perceptual salience.

Several studies have shown that the precision of numerosity discrimination can be predictive of current and/or future mathematical performance (Anobile et al., 2013b; Anobile et al., 2016; Chen and Li, 2014; Halberda et al., 2008; Libertus et al., 2011, 2013). At the lower end of the spectrum, some dyscalculic children have been shown to present abnormally high numerosity thresholds (Mazzocco et al., 2011; Piazza et al., 2010). Accordingly, one influential theory posits that numerosity representations are foundational for higher-level numerical skills and that impairments in these representations may prevent individuals from understanding the semantic meaning of symbolic numerals, and higher level arithmetic (Butterworth, 2005; Butterworth et al., 2011; Butterworth and Kovas, 2013; Dehaene et al., 2003; Landerl et al., 2004). However some authors observed slower and less accurate responses during digits, but not non-symbolic comparisons in children with mathematical learning disabilities, and proposed that the source of the difficulties was in linking number symbols to magnitude representations, rather than in numerosity processing per se (Rousselle and Noël, 2007).

Beyond these core deficit hypotheses, more comprehensive views explain the heterogeneity of dyscalculia and the normal development of different components of mathematical cognition by taking into account also domain general cognitive abilities, such as working memory, attention and inhibition (Cragg and Gilmore, 2014; Fias, 2016; Fias et al., 2013; Geary and Moore, 2016; Houdé and Tzourio-Mazoyer, 2003; Linzarini et al., 2015; Menon, 2016; Poirel et al., 2012; Vanbinst et al., 2014; Vanbinst and De Smedt, 2016).

In particular, recently it has been suggested that mathematical achievement could be more related to the ability of the subjects to inhibit responses to task-irrelevant features rather than to the numerosity acuity itself: Gilmore et al. (2013) found that in typically developing children the correlation between weber fraction and mathematical skills was significant only when other quantitative features varied incongruently with number, and that weber fractions were no longer predictive of calculation ability once separate measures of inhibitory skills were included. Similarly, the performance of dyscalculic children during non-symbolic numerical comparisons was reported to be particularly affected by the congruency with other visual perceptual cues, (Bugden and Ansari, 2016; Szűcs et al., 2013). On the basis of these findings it has been suggested that the previously described relation between numerosity discrimination and arithmetic performance across the general population, as well as the particularly impaired numerosity acuity in some dyscalculic subjects, would not be due to a dedicated enumeration capacity being foundational as commonly assumed, but to a more domain-general deficit in executive function and especially inhibitory skills, manifesting as a poor ability to discard task-irrelevant features during numerosity judgement

The aims of the work described in this manuscript were two-fold. First, in normal adult subjects, we wanted to determine what is the capacity of numerosity to interfere with the judgement of another quantitative dimension (average item size) and how it compares to the degree of interference of that feature onto numerosity under conditions of equated perceptual discriminability. We chose average item size as an intuitive feature which is considered an explicitly encoded visual dimension, as number and density (Ariely, 2001; Chong and Treisman, 2005, 2003; Corbett et al., 2012; Sweeny et al., 2015). As summarized previously, unequal discriminability can affect the degree and direction of interference and merely equating physical ratios across magnitudes does not necessarily capture subjects’ perceptual sensitivity. Therefore, to determine the intrinsic capacity for interference more unambiguously, in a pilot study we measured perceptual precision for both average item size and numerosity in normal subjects, which then allowed us to equate the difficulty of the two tasks on average across subjects. We asked participants to make comparative judgments over the same sets but on the basis of either of the two dimensions.

Second, to arbitrate between the hypotheses of impaired number acuity versus domain-general inhibition deficits in dyscalculia we tested a group of adult dyscalculics with our novel paradigm. Having access to adult dyscalculics allowed us to extensively test them with different tasks and a large number of trials, enabling robust and fine-grained psychophysical measures that are much harder to obtain in children. Comparing dyscalculic participants’ performance with an age and IQ matched control group on average item size, in addition to numerosity discrimination, allowed us to directly evaluate, for the first time, the hypothesis according to which dyscalculia is associated to a general deficit of inhibitory control. If dyscalculics suffered from a generalized inhibition impairment and no domain specific number sense deficit, we would expect them to present stronger interference than the control group irrespective of the task-relevant dimension (numerosity or average item size). On the contrary, if decreased precision and / or enhanced interference in the dyscalculic compared to the control group was found only during the numerosity task but not during the average size task, this would refute the domain-general view and be more compatible with a domain-specific deficit in numerosity representation.

## Methods

### Subjects

Fifteen adults without mathematical impairment and ten adults with mathematical impairment participated in the study. Contacts with math impaired subjects were provided by our speech therapist collaborator to whom participants referred during childhood or adult age for evaluation. To be included in the dyscalculic group participants were required to (a) have been diagnosed with dyscalculia by a neuropsychologist or speech therapist during childhood or have suffered from major difficulty with math since very early in school; (b) to claim that the math difficulty interfered with their everyday life and career choice; (c) present no neurological disorder; (d) have completed at least secondary level education.

Participants included in the control group were required to (a) have had no difficulty learning mathematic, reading, writing and orthography during school; (b) not have any neurological disorder; (c) have at least secondary level education.

All subjects underwent an extensive neuropsychological assessment where indices of verbal and non-verbal intelligence, verbal and visuospatial working memory, reading abilities, inhibitory skills and mathematical performance were measured, to objectify differences in mathematical abilities and compare performance of the groups across more general domains.

One subject who initially claimed not to have any mathematical difficulties was excluded from the experiment because his/her performance was more than 2 standard deviations below the group mean for both intelligence indices and for more than one test measuring different components of mathematical abilities. Therefore fourteen adults in the control group (C group, age 29±7) and ten adults in the dyscalculic group (D group, age 28±11) were included in the main experiment.

All participants signed the informed consent. This study was conducted in accordance with the Declaration of Helsinki and under the general ethics protocol covering human research at Neurospin (Gif-sur-Yvette, France). The study was reviewed and approved by an institutional review board (ethics committee) before the study began (it received authorization from the CPP IDF 7 number 15 007 on May the 28th 2015 and from the Agence du Médicament on February the 13th 2015).

### Neuropsychological Assessment

The neuropsychological evaluation started with an anamnestic interview where the compliance with the inclusion criteria was verified for each participant.

After the interview, all subjects underwent neuropsychological testing. As a measure of verbal and non-verbal IQ, we selected two representative subtests of the Wechsler Adult Intelligence Scale IV edition (WAIS-IV): similarities and matrix reasoning, respectively. Verbal working memory was evaluated by means of the digit span subtest from WAIS-IV, while visuospatial working memory was measured with the Corsi-Block Tapping test. Reading abilities were evaluated with the “Alouette”, one of the most widely-used reading tests in France (Lefavrais, 1967). This is a timed test that requires participants to read aloud a brief text composed of existing regular and irregular words, arranged in a grammatically plausible manner within the sentence, but conveying no clear meaning overall.

The Stroop-Victoria test adapted for francophone subjects (Bayard et al., 2009) was administered to measure inhibitory skills, selective attention and processing speed. Participants were required to spell aloud as quickly as possible the color of the ink of a series of filled circles, of a list of words (‘mais’, ‘pour’, ‘donc’, ‘quand’, meaning ‘but’, ‘for’, ‘so’, ‘when’) and of a list of color words (‘jaune’, ‘rouge’, ‘vert’, ‘bleu’, meaning ‘yellow’, ‘red’, ‘green’, ‘blue’). Importantly the color of the ink used for the color words was always incongruent with the meaning (for example ‘bleu’ written in red). The interference index is calculated by dividing the time necessary to perform the task with the color words by the time needed to name the color of circles.

Finally, to assess mathematical abilities, subjects were evaluated with parts of the French battery TEDI Math Grands (Noël and Grégoire, 2015). This battery includes computerized tests evaluating basic numerical abilities. Accuracy and reaction times were recorded while the subjects were: 1) estimating the number of briefly presented items within the subitizing range; 2) comparing two single-digit Arabic numerals; 3) mentally performing single-digit multiplications and subtractions. Additionally, all the subjects underwent two subtests taken from the Italian battery for developmental dyscalculia (BDE) specifically targeting understanding of the semantic meaning of numerals (Biancardi and Nicoletti, 2004). In the first subtest, the subjects were asked to choose the largest of three vertically arranged Arabic numerals (one to three digits), while in the second one the subjects had to correctly place an Arabic numeral (one to four digits) in one of the four possible positions along a number line. Both of these tests measure response accuracy and overall response speed and were chosen for targeting the understanding of numerals’ semantic associations. Moreover, these tests were found by previous studies to best correlate with numerosity discrimination thresholds, compared to tasks evaluating transcoding, memory and automatization of procedures (Anobile et al., 2013b; Anobile et al., 2016).

### Analysis

Referring to standardized norms for adults, we calculated standard scores for the IQ subtests, for the verbal (digit) and visuospatial working memory and for the Stroop test. For the reading test we analyzed the time (in seconds) needed to read the proposed text and the number of errors. For the TEDI-MATH we analyzed the number of items to which subjects correctly responded and, when measured, the reaction time (in ms) needed to respond. Because accuracy and reaction time can often inversely trade off with each other, we reduced the number of measures by calculating the inverse efficacy (IE) score (Collins et al., 2017). IE score is calculated by dividing, for each participant, the mean RT by the proportion of correct responses. Results from the multiplication and subtraction test in the TEDI math were averaged together and the IE score Calculation was computed from the collapsed measures. As the two BDE tests were addressing the same semantic component of numeracy, we reduced them to one single value by averaging their scores. Similarly to the other tests, the IE score was computed.

To evaluate differences across groups, we compared the dyscalculic and control group’s performance using independent sample t-tests. These tests were applied to either the standardized test scores described (for the IQ, memory and Stroop tests) or to the raw scores in the cases where the norms did not cover the adult age range (in the case of the math and reading tests). When Levene’s test was significant, the corrected value, not assuming the equality of variances, was reported.

## Psychophysical experiment

### Stimuli and procedures

Stimuli consisted in heterogeneous arrays of dots, half black and half white, briefly presented (200 ms) on a midgray background. Dots were constrained to be at least 0.25° apart from each other, to not overlap with the fixation point and to fall within a virtual circle of either 7.6° or 5.8° diameter of visual angle. Arrays of dots were designed to be sufficiently sparse to target the ‘number regime’ and to avoid the contribution of texture density processing mechanisms that might come into play when item segregation is not possible (Anobile et al., 2015, 2013a). Indeed, the largest number of dots displayed within the smallest total field area at the highest eccentricity yielded a density of 0.75 dot/deg^2^, therefore still falling within the number regime. The sets of dots generated were orthogonally varying in mean size and numerosity. In different sessions participants were asked to perform two different tasks. During the ‘numerosity task’ sessions subjects were asked to choose which one of two stimuli was more numerous, regardless of the mean size of the dots. During the ‘average size task’ sessions instead, subjects were asked to choose the array containing the dots with the largest average size. Results from a pilot study on eight subjects were used to estimate the just noticeable distance (JND) on a logarithmic scale for numerosity (0.15) and average size (0.08, when expressed as a function of average item diameter change, or 0.15, when expressed as a function of average item area change). Based on these measurements, we chose the ratios to be compared in each task to be adapted to each dimension’s JND. The unattended dimension was chosen to only take the most extreme values. In the set of stimuli used for the number discrimination task the arrays contained 5, 6, 8, 12, 17 and 20 dots (ratios 0.5, 0.6, 0.8, 1.2, 1.7, 2 with respect to the reference of 10 dots), and these dots could be presented with either small (0.25°) or large (0.5°) average diameter. The arrays used for mean size discrimination contained dots with average diameter of 0.25, 0.27, 0.3, 0.40, 0.46 and 0.5 visual degrees (ratios 0.71, 0.77, 0.86, 1.15, 1.3, 1.4 with respect to the reference of 0.35 visual degrees) presented with either few (5) or many (20) dots. This is equivalent to saying that, expressed in terms of average item area, we tested 0.05, 0.06, 0.07, 0.13, 0.16 and 0.19 visual square degrees,corresponding to the same ratios as those tested for numbers (0.5, 0.6, 0.8, 1.2, 1.7, 2). In both tasks, the test stimuli were compared to a reference stimulus containing 10 dots with 0.35° average item diameter (or 0.1 degree square of average item area) within the same total field area as the test stimulus.

For each array, single dots diameters were derived from a symmetric interval around the mean size, which was linearly subdivided into as many bins as the number of dots included in the array. To prevent arrays with larger mean sizes from subjectively appearing to be composed by less variable dot sizes than the smaller ones, as it was the case when using a constant interval across all sizes, we scaled the size of the interval with mean size. The intervals spanned ±0.09, ±0.11, ±12, ±0.15, ±0.17, and ±0.19 visual degrees around the respective mean size. Examples of the stimuli used in the two tasks are shown in Fig 1A.

**Fig 1.**
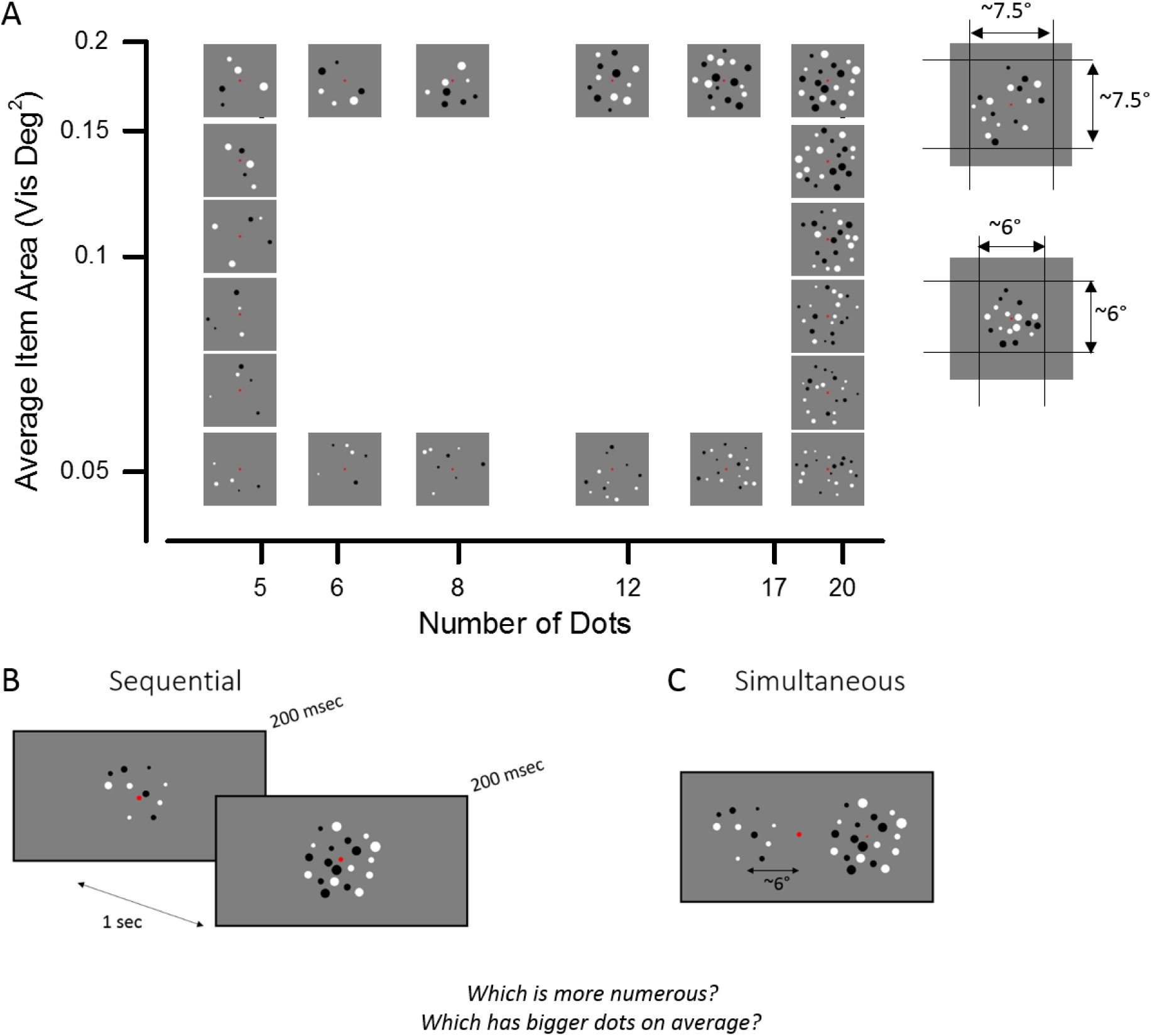
(A) Example of stimuli in the numerosity and average item size comparison tasks. The set of stimuli was created with two different total field areas of ∼7.5° and ∼6° diameter. (B, C) The two stimuli were shown either in sequential or simultaneous presentation mode. In separate sessions, participants were asked to judge which array contained more dots or which one contained the dots with the larger average size.

Visual stimuli were presented in a dimly lit room on a 14-inch HP screen monitor with 1024×768 resolution at refresh rate of 60 Hz, viewed binocularly from approximately 60 cm distance. Stimuli were generated and presented under Matlab 9.0 using PsychToolbox routines (Brainard, 1997).

The order of the two tasks was counter-balanced between subjects with half of the subjects starting with the numerosity task and the other half with the mean size task. In different days, the control group was tested with two experiments. The stimuli and tasks were the same in the two experiments, but in Experiment 1 the stimuli were presented sequentially, while in Experiment 2 they were presented simultaneously (Fig 1B). The order of the experiments, i.e. the order of presentation modes (sequential/simultaneous), was counter-balanced across subjects, with half of the subjects starting with Experiment 1 and the other half with Experiment 2. During the sequential presentation, the two patches were presented in the center of the screen one after the other, separated by a 1 s interval. When presented simultaneously, the two sets of dots appeared centered at 6 degrees of eccentricity along the horizontal meridian with respect to the central fixation point. Test and reference stimuli could appear either as first or as second stimulus during the sequential presentation and to the left or to the right of the fixation point during the simultaneous presentation. After stimulus presentation the subjects’ responses were recorded by button press. Subjects were instructed to press the left arrow to select the stimulus on the left or the first stimulus in the simultaneous and sequential presentation respectively, and to press the right arrow to select the right or the second stimulus.

In Experiment 3 we tested the dyscalculic group with the simultaneous presentation only, in order to minimize short-term memory load.

Each session started with instructions and 12 practice trials, after which the experiment started. Each subject performed three sessions of one task, followed by a pause and another three sessions of the other task. For each task each one of the 6 comparison ratios was presented 72 times: 2 unattended magnitudes (small and big during the number task and five or twenty dots during the size task), 2 possible total field areas, 2 possible spatial positions/presentation orders with respect to the reference (left-right/first-second) repeated 3 times in each one of the 3 sessions. A total of 432 trials per task were collected and used for the analysis in each experiment.

### Analysis

For each subject we quantified the effects of experimental manipulations on response accuracies as well as on parameters derived from fitting the psychometric functions.

To assess the effect of congruency across dimensions as well as the effect of ratios within dimension, we computed the proportion of errors as a function of the ratio of the attended dimension after splitting for congruency across dimensions. In the ‘congruent’ trials, the unattended dimensions varied in the same direction as the attended one with respect to the reference. On the contrary, in the ‘incongruent’ trials the attended and the unattended dimensions varied in opposite directions. For example, five small dots and twenty big dots were classified as ‘congruent’ trials, while five big dots and twenty small dots were classified as ‘incongruent’ trials. The congruency effect corresponds to more errors for the incongruent compared to congruent trials.

To quantify overall precision in both number and mean size judgments, we computed the just noticeable difference (JND) for each task, presentation mode and group. The percentage of test trials with “greater than reference” responses was plotted against the log-transformed difference between test and reference and fitted with a cumulative Gaussian function using Psignifit toolbox (https://github.com/wichmann-lab/psignifit). The 50% point estimated the point of subjective equality (PSE), and the difference between the 50% and the 75% points yields the just notable difference (JND).

A common way in psychophysics to measure interference is to estimate the response bias, quantified as the shift of the psychometric curve from the veridical value under different conditions, and allowing to appreciate the strength and direction (over vs underestimation) of the influence from the unattended dimension Therefore, to estimate the bias from the unattended dimension, we fitted the subjects’ responses after splitting the entire dataset for the different magnitudes (small or big) of the unattended dimension: during the mean size task, the ‘unattended small’ trials only included arrays containing five dots, while the ‘unattended big’ trials included only the twenty dot arrays. During the numerosity task, an equivalent subdivision was made based on small and large mean item size. A systematic shift of the PSE away from 0 as a function of unattended magnitude would suggest a bias from the unattended dimension. We calculated for each subject the signed difference between the two PSE estimates obtained when fitting the data after splitting for the magnitude of the unattended dimension (small-big). Moreover, since previous studies have shown that the direction of the bias from the unattended dimension is not necessarily the same for all subjects (DeWind et al., 2015) and this was also observed in our results, we computed in addition an unsigned bias, which measures the overall degree of interference effect irrespective of its direction, by taking the absolute value of the above described difference in PSE for small and large magnitude of the unattended dimension.

Effects of the experimental manipulations on the different measures described were tested statistically with repeated measures ANOVAs, including group as a between subject factor when comparing the control and dyscalculic group. In case of significant higher order interactions between factors, lower order interactions or main effects are not reported.. In case of significant interactions, post-hoc tests were always performed with adjustments for multiple comparisons (Bonferroni correction). One sample t-tests were used to test whether signed biases were significantly different from 0.

We further performed correlation analyses based on Pearson correlation, to test for a relation between the number and size bias with the subject’s sensitivity for these properties, as well as with the mathematical performance defined as IE calculation score, with and without regressing out the effect of group.

## Results

### Neuropsychological Assessment

The neuropsychological assessment verified the fulfillment of the inclusion criteria for all participants. Until recently, dyscalculia was a relatively unknown and underestimated disorder, therefore it is extremely rare to find adult dyscalculics with an established pre-existing diagnosis. Yet three of our subjects included in the dyscalculic group had been diagnosed with dyscalculia during childhood. None of the subjects had any neurological disorders and they all reported having had access to appropriate education during school-age. All the subjects had at least secondary level education.

Only subjects in the dyscalculic group claimed having had learning difficulties and major problems in acquiring mathematical skills since the early school years. Despite the fact that most of them (9 out of 10) had had intensive compensatory training and/or supporting private lessons, they all affirmed that their deficits continued to persist and to have an impact on their everyday life. Almost all of these subjects (8 out of 10) reported having at least one relative with difficulty in either mathematics, reading, writing or orthography. Four subjects in the dyscalculic and three subjects in the control group were born before the term (five subjects were born less than one month before the term, one subject in the control group two months before the term and one subject in the dyscalculic group four months preterm). Two subjects in each group were left handed.

The dyscalculic and control group did not significantly differ in age, verbal and non-verbal IQ, reading accuracy, verbal working memory and performance in the Color-Stroop test (all p-values>0.05, see Table 1 for descriptive statistics and tests across groups). The two groups significantly differed in reading speed (t(22)=2.24, p<0.05), visuo-spatial working memory (t(22)=-4.05; p<0.01), and basic numerical as well as arithmetic tests. In particular, dyscalculic and control group differed in accuracy in the subitizing task (t(22)= −2.61; p<0.01) and in IE scores for digit comparison (t(22)= 3.54; p<0.01), and calculation (t(22)= 2.30; p<0.05). Detailed results for RTs and accuracy during the individual tasks are listed in Table 1. Dyscalculics were significantly slower in digit comparison (t(22)= 3.30; p<0.01) and made more errors in mental multiplication and subtraction with respect to the control group (t(22)= - 4,74; p<0.01, t(22)= −2.83; p<0.01). Additionally IE score in the two subtests of the BDE battery differed across groups (t(22)= 4.40; p<0.01). Here dyscalculics were significantly less accurate and slower than participants in the control group (t(22)= −3.54; p<0.01; t(22)= 3.96; p<0.01).

**Table 1.**

|  | Control group<br>(N=14) | Dyscalculic group<br>(N=10) | Statistical<br>analysis |
| --- | --- | --- | --- |
|  | Mean (STD) | Mean (STD) | t-value |
| Age | 29 (7) | 28 (11) | -0.26 |
| IQ |  |  |  |
| Similarities | 12 (3) | 12 (3) | -0.56 |
| Matrices | 10 (2) | 9 (2) | -1.63 |
| Reading Ability |  |  |  |
| Time (seconds) | 98 (19) | 121 (32) | 2.24* |
| N errors | 4 (3) | 6 (6) | 0.91 |
| Working memory |  |  |  |
| Verbal | 11 (3) | 9 (2) | -1.94 |
| Visuospatial | 13 (2) | 9 (2) | -4.05** |
| Color Stroop |  |  |  |
| Inhibition Index | 12 (2) | 13 (2) | 1.26 |
| Arithmetical tests |  |  |  |
| TEDI – MATH (no of items) |  |  |  |
| Subitizing (36) | 34 (2) | 30 (4) | -2.61 ** |
| Digit Comparison |  |  |  |
| Accuracy (48) | 46 (1) | 46 (2) | -0.11 |
| Reaction Time (ms) | 598 (67) | 763 (147) | 3.30 ** |
| IE score Digit | 6 (0.6) | 7.8 (1.4) | 3.54 ** |
| Multiplication |  |  |  |
| Accuracy (20) | 17 (2) | 13 (2) | -4.74 ** |
| Reaction Time (ms) | 1913 (497) | 2961 (1723) | 1.86 |
| Subtraction |  |  |  |
| Accuracy (20) | 19 (1) | 17 (2) | -2.83** |
| Reaction Time (ms) | 1797 (734) | 2946 (2095) | 1.66 |
| Calculation (x and -) |  |  |  |
| IE score Calculation | 41 (14) | 79 (50) | 2.30 * |
| BDE |  |  |  |
| Accuracy (34) | 34 (0.4) | 32 (1.2) | -3.54 ** |
| Reaction Time (s) | 71 (12) | 114 (31) | 3.96 ** |
| IE score BDE | 0.3 (0.06) | 0.6 (0.16) | 4.40 ** |
| DD differs significantly from controls at: |  |  |  |
| * p=0.05 |  |  |  |
| * at p<0.05 |  |  |  |
| **At p<0.01 |  |  |  |

### Experiment 1: sequential judgments in subjects without math difficulty

In Experiment 1, participants performed the numerosity and average item size task with sets of dots presented sequentially. Weber fractions were in line with those expected based on the pilot study measurements, being equal to 0.16±0.08 for number and to 0.16±0.03 for mean size judgments. To evaluate whether participant’s responses were affected by changes in the unattended dimension we compared the proportion of errors in congruent versus incongruent trials and the PSE values obtained by fitting psychometric curves after separating the trials according to the magnitude of the unattended dimension.

#### Congruency effect

Fig 2 illustrates the proportion of errors made in the numerosity (Fig 2A) and average size (Fig 2B) task when judging congruent (solid lines) and incongruent (dashed lines) trials as a function of the ratio tested (grouped in far, medium and close with respect to the reference, as symmetric values were tested). As expected, in both tasks subjects made on average more errors when judging the most difficult ratios. Interestingly, numerosity judgments were not affected by congruency, while the proportion of errors made during the average size task was higher for the incongruent trials with respect to the congruent ones. The congruency effect observed in the average size task was smallest for the easiest ratios and tended to increase as the distance between test and reference decreased.

**Fig 2.**
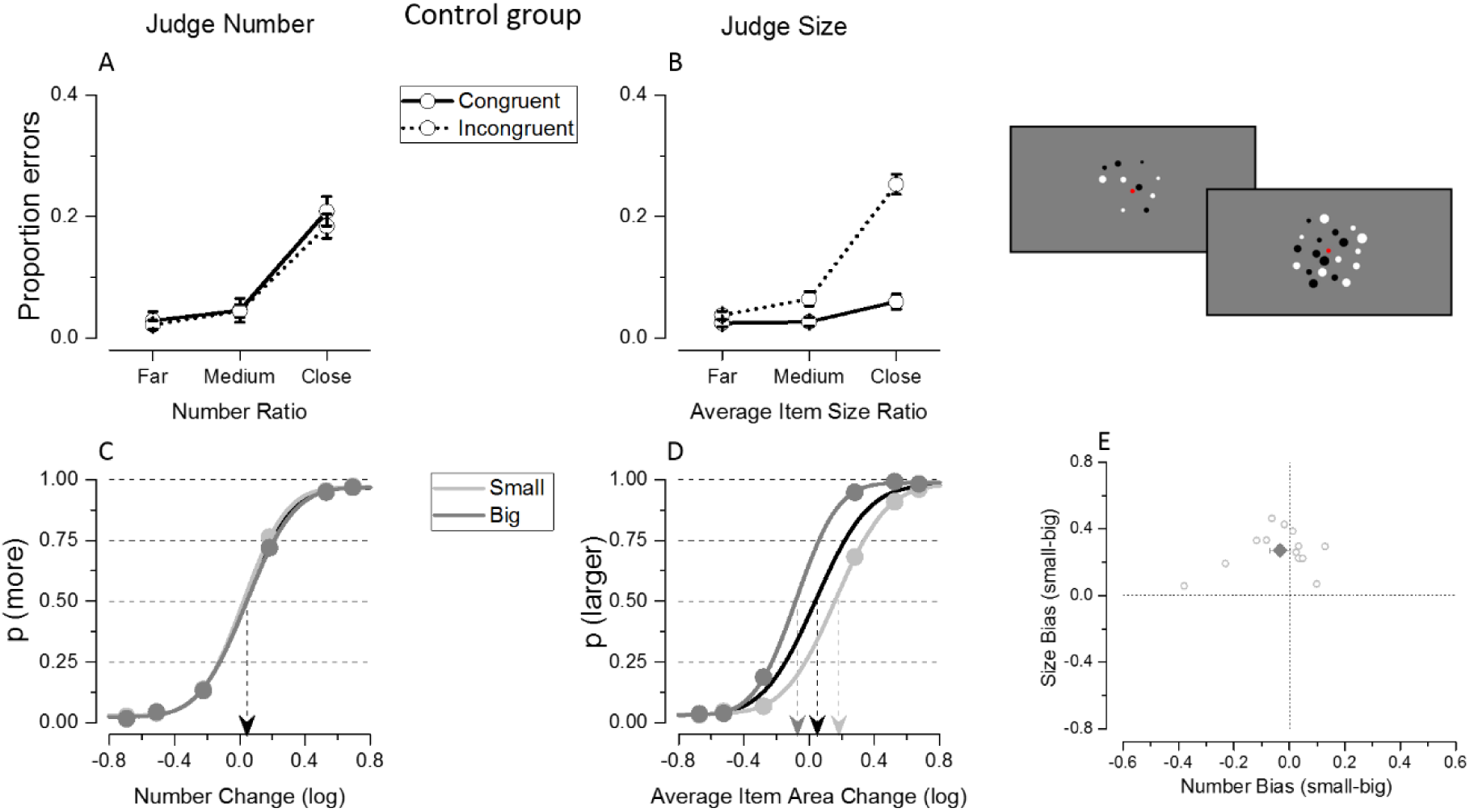
Results from Experiment 1 where control subjects were tested with sequentially presented stimuli. (A-B) Proportion of errors as a function of ratio of the attended dimension during numerical (A) and average size (B) judgments. Different lines show the error rate when participants were tested with congruent (solid line) or incongruent (dotted line) trials. (C-D) Psychometric functions for the control group for the number (C) and average size (D) tasks. Black curves fit the entire dataset while light and dark gray curves fit trials that are the small and the big, respectively, within the unattended dimension. Data in E show the average (dark big diamond) and single subjects’ PSE difference (light gray small circles) during numerosity (on the x axis) and average size (on the y axis) comparison when the dataset was split for the magnitude of the unattended dimension (small-big).

To quantify these effects the proportion of errors was entered in a 2 (task: judge number/mean size) x 2 (congruency: congruent/incongruent) x 3 (ratios) repeated measure ANOVA. The significant triple interaction between task, congruency and ratio (F(2,26)=21.94; p<10^−5^) and the post-hoc comparison tests confirmed that congruency affected accuracy differently during the two tasks as a function of the ratios to be compared. The congruency with the unattended dimension did not affect the proportion of errors made during the numerosity comparisons at any ratio tested (ratio far: p=0.45; ratio medium: p=0.95; ratio close: p=0.47). On the contrary, in the average size task the error rate during incongruent trials was smallest for the easier ratios and tended to increase as the comparison between arrays of different average sizes became more difficult (ratio far: p=0.12; ratio medium: p=0.002; ratio close: p<10^−5^).

#### Interference from the unattended dimension

To test whether and in which direction the unattended magnitude was biasing participants’ responses, we evaluated the shift along the x axis of the psychometric curves when fitted using trials where the unattended dimension was small or big. As shown in Fig 2C, the two curves overlapped when fitted on the average of participants’ numerical judgments, indicating the absence of bias. On the other hand, the two average psychometric functions clearly separated when fitted on the average size responses (Fig 2D), suggesting that in this case participants were systematically influenced by the magnitude of the unattended dimension, i.e. the numerosity of the patch. Specifically, participants tended to overestimate average size when presented with large numerosity (dark gray curve shifted towards the left on the x axis) and to underestimate it when presented with small numerosity (light gray curve shifted towards the right on the x axis). In line with these observations, the 2 (task: judge number/mean size) x 2 (unattended magnitude: small/big) repeated measure ANOVA performed on PSEs estimates showed a highly significant interaction between task and magnitude of the unattended dimension (F(1,13)=52.17, p<10^−5^), with PSE estimates differing between small and large unattended magnitude only for the average size task (p<10^−5^) but not for the numerosity task (p=0.37).

The absence of a group average bias when judging numerosity might have been potentially due to strong but opposite sign effects at the single subject level which cancelled each other out. However this was not the case, as illustrated by the single subjects’ differences in PSEs estimates (small-big) when judging number in Fig 2E: all subjects’ signed biases were clustered very closely around zero, leading to an overall PSE difference that was not significantly different from zero (t(13)=-0.91, p=0.37). The PSE shift due to numerosity interference affecting average size judgments was systematically occurring in the same direction across subjects and was significantly different from zero (t(13)=8.53, p<10^−5^).

### Experiment 2: simultaneous judgments in subjects without math difficulty

To assess whether potential differences in attentional or working memory load due to different presentation modes modulated the interference effect, in Experiment 2 participants were tested with the numerosity and average size tasks, but with stimuli presented simultaneously in the periphery instead of sequentially in the center of the screen. Average Weber fractions were 0.17±0.03 for number judgment and 0.2±0.05 for mean size judgments, therefore similar to the ones obtained in the previous experiment, but slightly higher probably due to the peripheral presentation of the stimuli. Interference from the unattended dimension was evaluated by applying the same analysis and statistical tests as used in Experiment 1.

#### Congruency effect

The proportion of errors was entered in a 2 (task: judge number/mean size) x 2 (congruency: congruent/incongruent) x 3 (ratios) repeated measure ANOVA. When stimuli were simultaneously presented, similarly to what was observed with sequential displays, the triple interaction between task, congruency and ratio (F(2,26)=16.76, p<0. 10^−5^) was significant. Numerical judgments were never affected by changes in the unattended dimension (ratio far: p=0.82; ratio medium: p=0.48; ratio close: p=0.22), while congruency modulated the average proportion of errors made during the average size task, with the effect being stronger as the ratios to compare became more difficult (ratio far: p=0.003; ratio medium: p=0.002; ratio close: p<0. 10^−5^, Fig 3 A and B).

**Fig 3.**
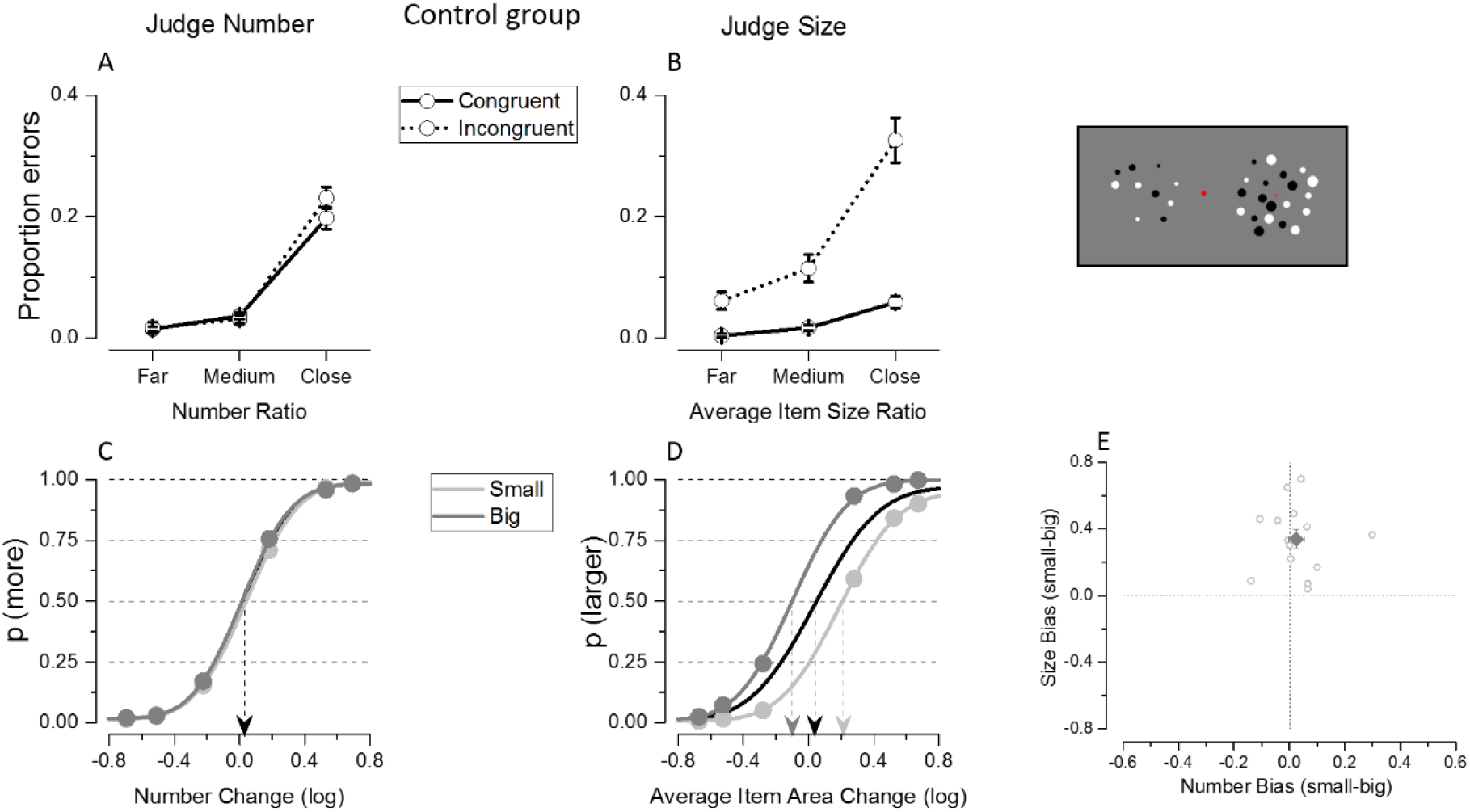
Results of Experiment 2 where the control group was tested with simultaneous presentation. Results show a similar pattern despite the change in presentation mode. Congruency effect and bias from the unattended dimension are evident in the proportion of errors and group average fits during the average size task, but not during the numerosity task.

#### Interference from the unattended dimension

When stimuli were presented simultaneously, the irrelevant dimension interfered with participant’s judgments in a way very similarly to when they were shown sequentially. Indeed while participant’s judgments did not differ based on the magnitude of the unattended dimension when judging numbers, they tended to over- (under-) estimate sizes when presented with large (small) numerosity (Figs 3 C and D). A 2 (task: judge number/mean size) x 2 (unattended magnitude: small/big) repeated measure ANOVA was performed on PSE estimates. The significant interaction between task and magnitude of the unattended dimension (F(1,13)=25.26, p<10^−5^), confirmed that PSE estimates did not differ during numerosity judgments (p=0.37), while they were significantly different when participants were comparing average sizes (p<10^−5^). When judging numerosity, most of the subjects’ differences in PSE estimates were clustered around zero, and as a consequence of this the bias was not significantly different from zero across subjects (t(13)=0.92, p=0.37). On the other hand the unattended number of dots systematically biased average size judgments in the same direction across subjects, leading to a significant difference from zero (t(13)=6.16, p<10-5; Fig 3E).

### Comparison between simultaneous and sequential judgments in subjects without math difficulty

In the control group, weber fractions were on average slightly higher when stimuli were presented simultaneously than when they were presented sequentially (w-values for numerical judgment simultaneous vs sequential: 0.17±0.03 vs 0.16±0.08; w-values for average size judgment simultaneous vs sequential: 0.20±0.05 vs 0.16±0.03). However, presentation mode did not significantly impact on precision for both visual dimensions (no significant main effect of presentation mode: F(1,13)=1,97; p=0.18; no significant interaction between task and presentation mode F(1,13)=1,17; p=0.29).

To evaluate whether the different attentional and working memory load recruited when presenting stimuli simultaneously or sequentially modulated the strength of interference from the unattended dimension, the proportion of errors and PSE biases measured in Experiment 1 and 2 were directly compared.

The proportion of errors was entered in a 2 (presentation mode: sequential/simultaneous) x 2 (task: judge number/mean size) x 2 (congruency: congruent/incongruent) x 3 (ratios) repeated measure ANOVA. The significant triple interaction between task, congruency and ratio (F(2,26)=42,07; p<10^−5^) showed that, independently from the presentation mode, congruency significantly modulated error rate during average size (ratio far: p=0.002; ratio medium: p=0.001; ratio close: p<10^−5^), but not during numerical judgments (ratio far: p=0.28; ratio medium: p=0.62; ratio close: p=0.79). Interactions between the presentation mode and the other factors were not significant, suggesting that different presentation modes did not change the results (interaction between presentation mode, task and congruency: F(1,13)=1.30; p=0.27; interaction between presentation mode, task and ratio: F(2,26)=0.68; p=0.51; interaction between presentation mode, congruency and ratio: F(2,26)=1.83; p=0.17; interaction between presentation mode, task, congruency, and ratio: F(2,26)=0.53; p=0.59).

A 2 (presentation mode: simultaneous or sequential) x 2 (task: judge number/mean size) x 2 (unattended magnitude: small/big) repeated measure ANOVA was performed on PSE values. The significant interaction between task and magnitude of the unattended dimension (F(1,13)=64.31; p<10^−5^) showed that, independently from the presentation mode, the PSEs estimates were affected by the magnitude of the unattended dimension only during the average size (p<10^−5^), but not during the numerosity comparisons (p=0.79). Moreover also the interaction between task and presentation mode was significant (F(1,13)=5.96; p=0.03), with PSEs for average size being overall slightly larger during simultaneous with respect to sequential presentation (p=0.01), while no presentation mode related difference was observed in the overall PSEs estimates during numerical judgments (p=0.30). Other interactions between presentation mode and the other factors were not significant showing that the different presentation modes did not alter the strength of the bias from the unattended dimension (interaction between presentation mode and magnitude of the unattended dimension: F(1,13)=1,87; p=0.19; interaction between presentation mode, task and magnitude of the unattended dimension: F(1,13)=0.02; p=0.89).

To sum up, in the group of adult subjects without math difficulties, incongruent information from the unattended dimension increased the proportion of errors only when participants were comparing average size, but not when they were comparing numerosity. The congruency effect observed in the average size task was particularly strong when difficult ratios were tested and it was smaller for the easiest comparisons. The magnitude of the unattended dimension biased participants’ responses so that they judged more (less) numerous arrays as containing larger (smaller) average sizes. On the other hand, the magnitude of the irrelevant information did not bias numerical judgments. Differences in attentional and working memory recruitment caused by simultaneous or sequential presentation of the stimuli did not affect these results.

### Experiment 3: simultaneous judgments in the dyscalculic group

In Experiment 3, a group of adult dyscalculic subjects performed the numerosity and average size tasks with stimuli presented simultaneously as in Experiment 2. Weber fractions were equal to 0.21±0.07 for numerical judgments and to 0.23±0.05 for mean size comparisons. To evaluate the interference from the unattended dimension during both tasks, the same analysis and statistical tests as used in the previous experiments were applied.

#### Congruency effect

Differently from what was observed in the control group, in the dyscalculic group the congruency with the unattended dimension affected both numerosity and size comparisons (Figs 4 A and B). Indeed the proportion of errors made during the numerosity task was on average higher for the incongruent trials with respect to the congruent ones, as it was the case for the average size task.

**Fig 4.**
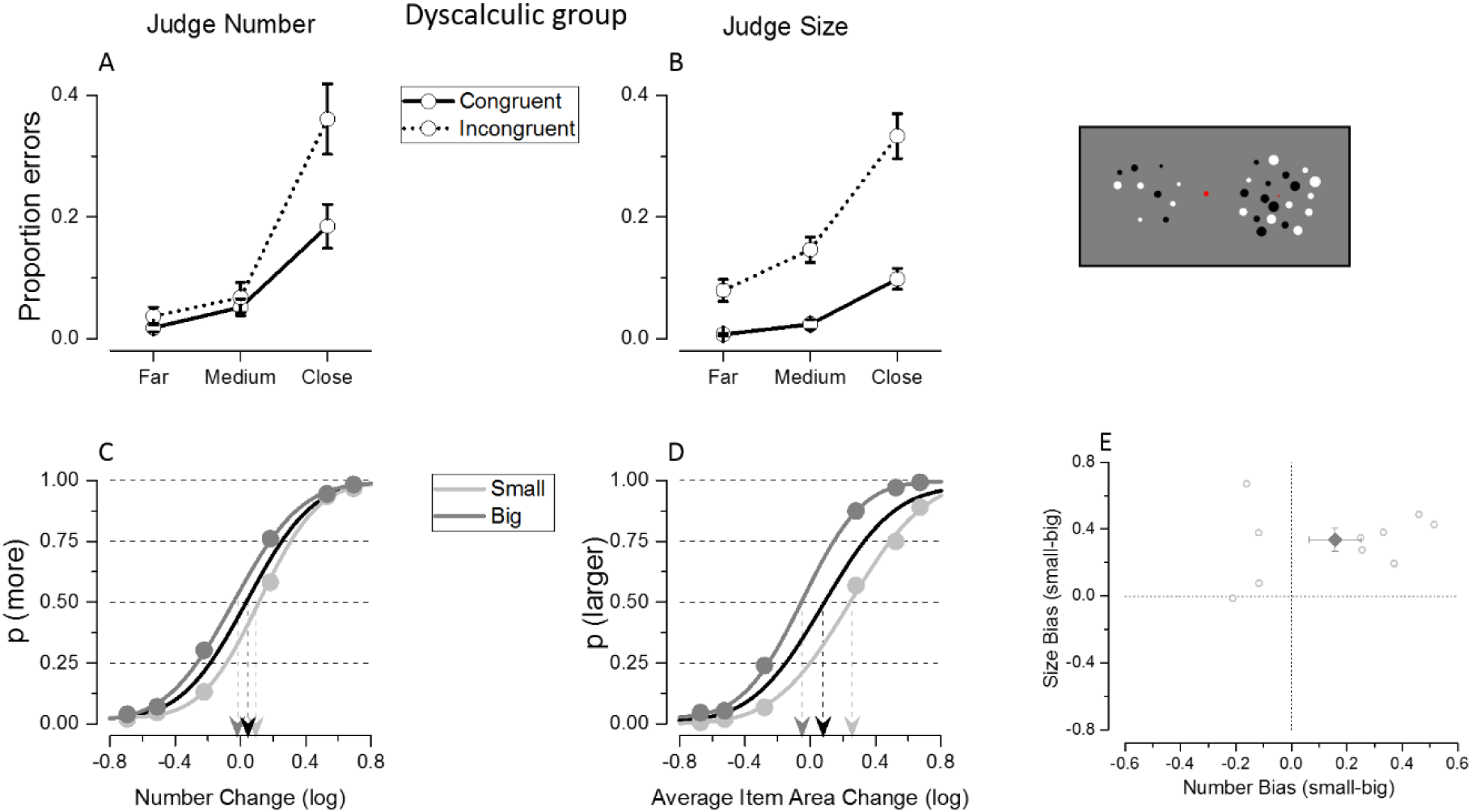
Results of Experiment 3 where the dyscalculic group was tested with simultaneous presentation. Differently from the control group (Fig 3), a tendency for congruency effects in accuracy at the most difficult numerical ratios, and bias from the unattended dimension in the group average fits, are visible during numerosity judgments.

These effects were quantified by entering the proportion of errors in a 2 (task: judge number/mean size) x 2 (congruency: congruent/incongruent) x 3 (ratios) repeated measure ANOVA. The interaction between task and ratio was significant (F(2,18)=12.05; p<10^−5^) because, independently from the congruency, the overall error rate during numerical judgments for the most difficult ratios was higher than the one during the average size task for the same ratio (p=0.03). The interaction between congruency and ratio was significant (F(2,18)=8.7; p=0.002), and equal in the two tasks (interaction between task, congruency and ratio: F(2,18)=0.34; p=0.71).This is because the judgments during the numerosity and average size task were both affected by congruency (interaction between congruency and task: F(1,9)=2.59; p=0.14) which was similarly affecting all the ratios tested: in both tasks, the strength of the congruency effect was smaller for the easier comparisons and tended to increase for the most difficult ratio tested.

#### Interference from the unattended dimension

Numerosity judgments of dyscalculic participants appeared to be biased by the magnitude of the unattended dimension, as shown in Fig 4C. Indeed, on average, they tended to overestimate numerosity when presented with large average item sizes and to underestimate it when shown with small average item sizes. The same tendency as the one observed in control subjects was found for the average size task, overestimating mean sizes when presented with higher numerosity and vice versa (Fig 4 D). Because both numerical and average size judgments were affected by the magnitude of the unattended dimension, in a 2 (task: judge number/mean size) x 2 (magnitude unattended: small/big) repeated measures ANOVA performed on PSE estimates in the dyscalculic group, the interaction between task and magnitude of the unattended dimension was not significant (F(1,9)=3.09; p=0.11). There was a significant main effect of the magnitude of the unattended dimension (F(1,9)=15.63; p=0.003), reflected by the psychometric curve’s shift and different PSE estimates. There was also a main effect of the task (F(1,9)=23.90; p=0.001) because the overall PSE estimated during the average size task was larger than the one during the numerosity task. However, while the magnitude of the unattended dimension showed a tendency to interfere with judgments in both tasks, post-hoc comparison showed that the PSE shift was significant only in the average size (p=0.001) and not in the numerosity task (p=0.10). Indeed, despite the fact that most subjects in the dyscalculic group showed stronger PSE shifts due to size interference during numerical judgments with respect to controls, the direction of the bias was not the same for all subjects: some dyscalculic subjects tended to strongly overestimate numerosity when presented with large average sizes, while some others tended to underestimate it (Fig 4E). Due to this fact, the overall effect tended to cancel out and the signed PSE bias (small-big), was not significantly different from zero for the number task (t(9)=1.78, p=0.10). On the other hand, the unattended numerosity significantly biased average size judgments (t(9)=5.10, p=0.01), in the same direction as the one shown by the control group.

### Comparison of control and dyscalculic groups

#### Overall Weber fractions

When judging average size the overall weber fraction was comparable between the dyscalculic and control subjects (w-values for dyscalculics vs controls: 0.23±0.05 vs 0.20±0.05). Compared to average size, the average between groups difference in numerical precision was larger (w-values for dyscalculics vs controls: 0.21±0.07 vs 0.17±0.03), with higher weber fraction for the dyscalculic group corresponding to an effect size (Cohen’s d) of 0.83 which suggests a relatively large difference in numerical precision between the two groups, yet not reaching statistical significance (no interaction between task and group F(1,22)=0.16; p=0.69).

#### Congruency effects in accuracy and signed biases

To evaluate whether the dyscalculic group’s judgments were differently affected by the irrelevant dimension with respect to the control group, we directly compared the proportion of errors and PSE values measured in the two groups when the same paradigm was used (i.e. when stimuli were simultaneously presented in Experiment 2 and 3).

The proportion of errors made by dyscalculic participants was compared to that of the control group by means of a 2 (task: judge number/mean size) x 2 (congruency: congruent/incongruent) x 3 (ratios) repeated measure ANOVA with group as between subjects factor. There appeared a significant quadruple interaction between task, congruency, ratio and group (F(2,44)=4.15, p=0.02) and the post hoc tests showed that with respect to the control group, the dyscalculic group made significantly more errors during the numerical task, when comparing the most difficult ratio of incongruent trials (differences across groups: ratio far: p=0.25; ratio medium: p=0.13; ratio close: p<0.03). Dyscalculics scored almost twice the errors made by the control subjects when presented with incongruent trials and difficult ratio (0.23±0.03 in controls vs 0.36±0.042 in dyscalculics). Both groups were equally affected by congruency during average size judgments and the congruency effect was not significantly stronger for the dyscalculic group with respect to the control group at any ratio tested (p>0.05 for all comparisons). The interactions between group and the other factors were not significant (interaction between task, congruency and group: F(1,22)=1,27, p=0.27; interaction between task, ratio and group: F(2,44)=1.33, p=0.27; interaction between congruency, ratio and group: (F(2,44)=0.56, p=0.57).

To evaluate group differences in signed bias a 2 (task: judge number/mean size) x 2 (magnitude unattended: small/big) repeated measures ANOVA was performed on PSE estimates with group as between subjects factor. As described earlier, the magnitude of the unattended dimension induced a bias in the dyscalculic group not only during average size comparisons, as in the control group, but also during numerosity judgments. When directly comparing the PSE bias across the dyscalculic and controls groups, the interactions between group and the other factor were not significant (interaction between task and group: F(1,22)=2.91; p=0.09; interaction between magnitude of the unattended dimension and group: F(1,22)=0.85; p=0.36; interaction between task, magnitude of the unattended dimension and group: F(1,22)=1.91; p=0.18). However, it is important to note that the absence of group differences in the bias induced by the unattended magnitude during numerical judgments could be explained by strong biases in opposite directions at the single-subject level in the dyscalculic group, resulting in only a modest signed PSE bias at the group level. On the contrary, the absence of group differences in the bias elicited by the unattended numerical magnitude during average size comparisons suggests that dyscalculics were not more affected by the unattended dimension with respect to the control group, given that the single subject’s signed bias was always in the same direction in both groups.

In sum, with respect to the control group, the dyscalculic group made more errors when asked to compare numerosity, although this was significant only for incongruent trials at the most difficult ratios. The congruency effect equally affected error rate across the two groups during the average size task. No significant difference was observed in the signed PSE biases across groups. This is likely a consequence of the fact that these measures are insufficiently representing the pattern present in the data, where in the dyscalculic group relatively strong biases are found but in opposite directions across different participants.

#### Unsigned bias

To evaluate whether the dyscalculic group showed an overall stronger interference (irrespective of its directions) from the unattended dimension with respect to the control group, the unsigned PSE biases measured during simultaneous judgment in Experiments 2 and 3 were directly compared. The dyscalculic group showed a much larger absolute bias mainly when judging numerosity, while the absolute size of interference was comparable across the two groups in the average size task (Fig 5). Accordingly, a one-way ANOVA (task: judge number/mean size) with group as between-subjects factor performed on the absolute biases yielded a significant interaction between task and group (F(1,22)=5.8; p=0.02). The additional post-hoc tests confirmed that, while dyscalculics’ numerical judgments were subject to a larger absolute bias with respect to the control group (p<10^−5^), for the average size task the groups did not differ significantly in the same measure (p=0.87). Thus, the dyscalculic group differed from the control group in the absolute degree of the interference, but crucially, this was only observed during numerosity, but not during size judgment.

**Fig 5.**
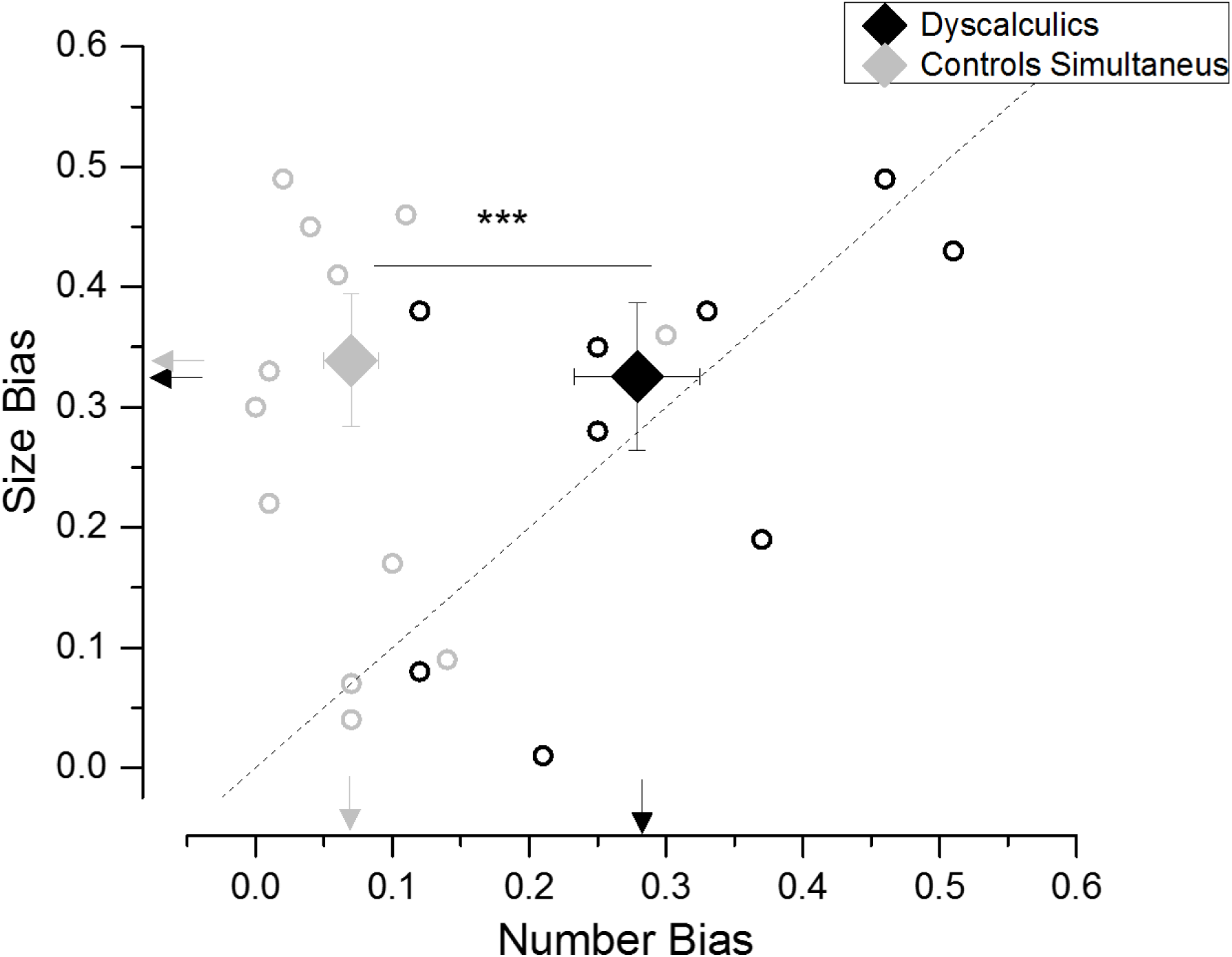
Absolute size of interference effect from the unattended dimension (unsigned PSE bias) arising when subjects in the control (gray symbols) and dyscalculic (black symbols) group judged numerosity (x axis) or average item size (y axis). Small circles represent individual subjects’ biases, large diamonds represent the group average ± sem. Arrows refer to average data values.

In sum, while participants in the control group could compare numerosity without a major influence from the unattended dimension, judging numerosity was more challenging for dyscalculic participants, and affected by the magnitude of the unattended size dimension, though not in the same direction across all participants. When asked to compare average sizes, dyscalculic participants were not more influenced by the numerical, irrelevant, information with respect to control participants, and the interference in this task was comparable across groups.

### Correlation analyses

To evaluate whether our data support the link between mathematical performance and precision of numerosity discrimination, we correlated the overall JND during numerical judgments and the IE score for mental calculation. We observed a significant correlation between mental calculation abilities and overall precision during numerical discrimination (r=0.6, p=0.002), even after controlling for group and inhibitory skills as measured by the color-word Stoop task (r=0.53, p=0.01). No significant correlation emerged when correlating mental calculation and overall precision during average size comparisons (r=0.32, p=0.12).

Under the hypothesis that stronger interference from the unattended dimension might emerge whenever the task difficulty increases, correlation analysis was performed to test whether the less precise subjects were also those whose judgment was more biased. To this aim we correlated the absolute magnitude of the bias with the overall JND. The numerosity interference during average size discrimination strongly correlated with overall precision in the average size task (r=0.71, p=0.0001), suggesting that as the difficulty of size discrimination increased (across subjects), interference from the unattended number dimension also increased (Fig 6A). Also, the correlation between average size interference during numerical judgments and JNDs for numerosity discrimination was significant (r=0.54, p=0.006, Fig 6B), however this was mainly due to the strong difference between groups.Correlations within individual groups did not reach significance, probably due to the small sample size available. Hence these correlations confirmed that less precise subjects were more influenced by the magnitude of the unattended dimension.

**Fig 6.**
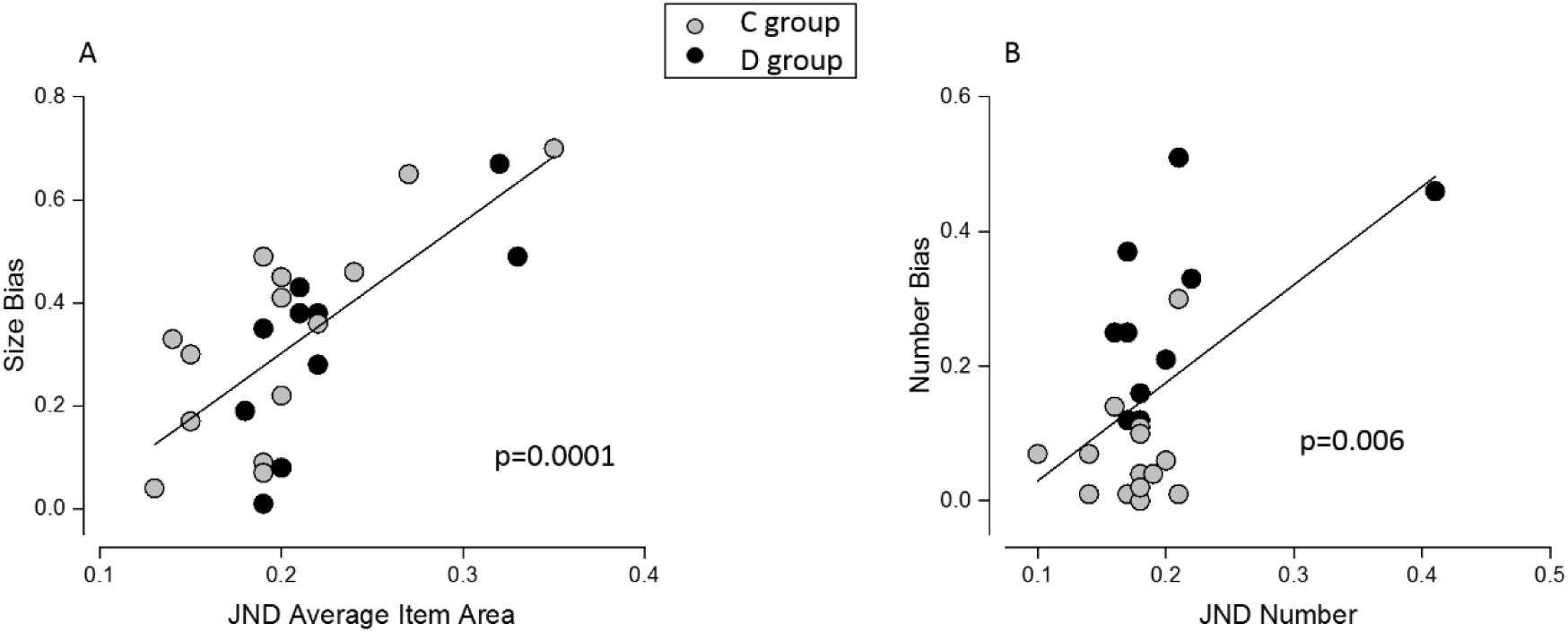
Correlation between the unsigned PSE bias and the overall precision during average size (A) and numerosity (B) judgments. Gray and black circles represents participants of the control and dyscalculic group, respectively.

To evaluate whether interference during the number and/or size task was related to mathematical performance, we correlated the absolute magnitude of the bias with the IE score for mental calculation. Numerical interference during average size judgments did not correlate with math performance (r=-0.02, p=0.90, Fig 7A). Instead, size interference during numerosity judgments highly correlated with mental calculation skills (r=0.60, p=0.002, Fig 7B), and this relation remained significant even when partialling out the group factor (r=0.41, p=0.04), the inhibitory skills as measured with the color-word Stroop task (r=0.63; p=0.001) and both group and inhibitory skills at the same time (r=0.47, p=0.03). Therefore the magnitude of the bias was related to mathematical ability only for numerosity, and not for size judgement. The subjects more proficient in mental calculation were also those who more efficiently discarded the irrelevant size information when comparing numerosity, while no relation was found with the bias during the average size task.

**Fig 7.**
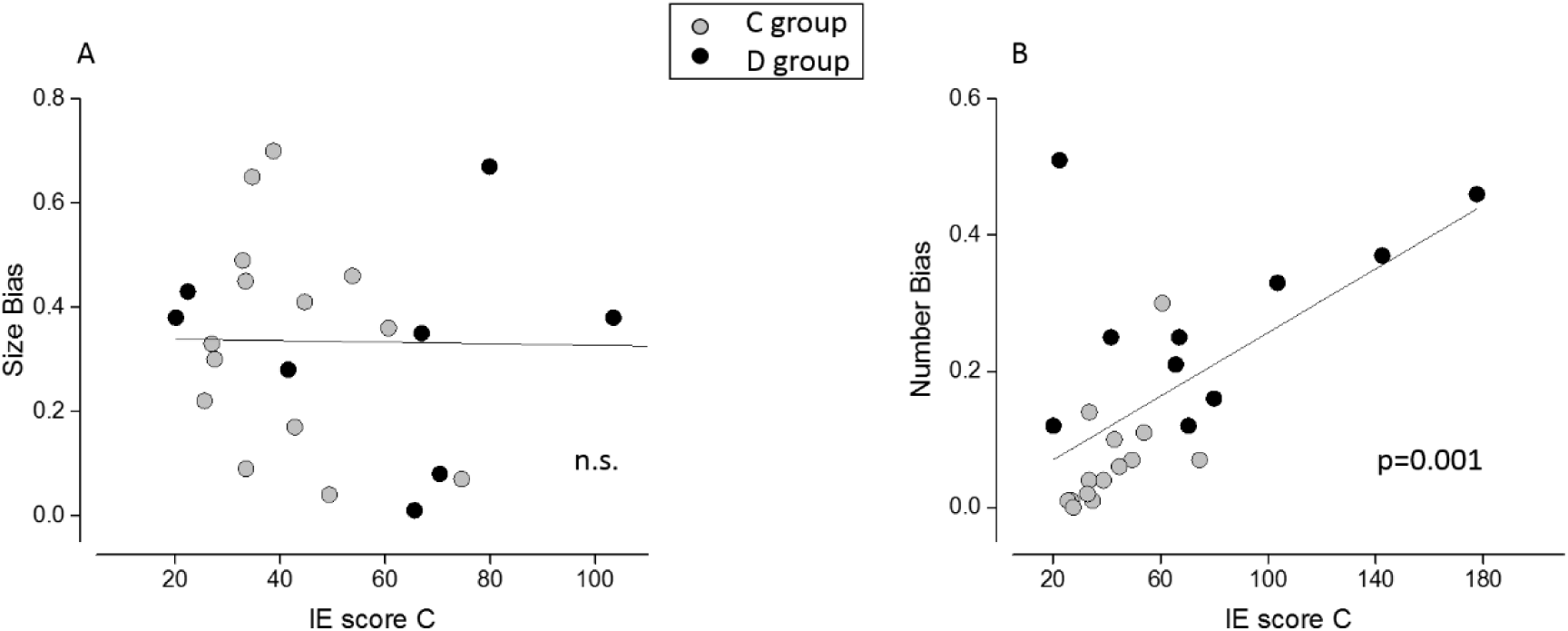
Correlation between the unsigned PSE bias in the average size (A) and numerosity task (B) and mental calculation skills. Only size interference during numerical judgment significantly correlates with math abilities, even when the factors of group and inhibitory skills are partialled out.

## Discussion

With the current study we aimed to evaluate for the first time the reciprocal interference between numerosity and another continuous dimension, average item size, under conditions where the perceptual discriminability was matched across tasks requiring judgement of one or the other dimension. Secondly, by testing dyscalculic adults on different quantitative dimensions of the same stimuli, we were able to directly compare the number sense deficit hypothesis of dyscalculia against the hypothesis of a domain-general inhibition deficit. Specifically, we evaluated whether dyscalculics were overall more subject to interference, in line with a general weakness in inhibiting task-irrelevant information, or whether numerosity judgment was preferentially affected by the unattended dimension, supporting a (domain specific) number sense deficit.

While participants without math impairments were able to compare numerosity without notable interference from the unattended dimension, they tended to overestimate mean sizes when presented with large numerosity, and tended to underestimate them when shown with small numerosity. This pattern of results was not affected by the presentation mode (sequential or simultaneous), suggesting that the interference pattern is unaffected by different allocation of attention or visuo-spatial memory load, at least as far as they relate to differences in presentation modes. Contrary to the controls, the dyscalculic group was strongly affected by the congruency of the irrelevant size information during numerosity judgment, although during average size judgement both groups were affected by the number of dots in the arrays to the same degree. Interestingly, only the ability to discard the irrelevant size information when comparing numerosity (but not vice versa) significantly predicted calculation ability.

The absence of interference from the unattended size dimension during numerosity judgement found in the present experiment in normal subjects contrasts with the often strong interference effects reported in the literature (Dakin et al., 2011; Gebuis et al., 2009; Gebuis and Reynvoet, 2012a, 2012b; Hurewitz et al., 2006; Leibovich et al., 2016a) even though in a few other cases, interference on numerosity judgement was also reported to be absent (Barth, 2008; Tokita and Ishiguchi, 2010). These differences may be due to a combination of several factors: our study used less difficult numerical ratios than some other studies, in combination with a relatively less extreme variation in the unattended dimension (DeWind et al., 2015; Hurewitz et al., 2006; Nys and Content, 2012; Tokita and Ishiguchi, 2010). To our knowledge, the present experiment is the first one to use stimuli that were calibrated based on previously measured thresholds for each dimension.

In addition, our study used relatively small numbers of items, contrasting with the much larger numerosities employed in some other studies (Bell et al., 2015; Dakin et al., 2011; Nys and Content, 2012). Behavioral evidence (Anobile et al., 2015, 2013a) supports a transition between a “number” and a “density” regime governed by different psychophysical laws. As a consequence, perceptual sensitivity for large numbers of densely spaced items can be predicted by the combined sensitivity to density and field area, but sensitivity for smaller numbers of well-segregated items cannot. For not too large numbers and not too densely spaced items, numerosity has also been shown to be the dimension that spontaneously drives humans’ and monkeys’ choices during quantity discrimination tasks (Cicchini et al., 2016; Ferrigno et al., 2017). Since our stimuli were explicitly chosen to fall into the “number” regime, they are more likely to have recruited processing mechanisms based on segmented items rather than indirect proxies to these such as the combination of texture density and area, which may have come into play in other studies. Of interest, Tokita and Ishiguchi (2010) already observed that the strength of size interference during numerosity judgments increased with numerosity, thus becoming stronger as stimuli were increasingly likely to move into the density regime. However, when testing smaller numbers of items, no interference emerged.

On the basis of the findings of Algom et al. (1996) in the number-size interference with numerical symbols we would have expected our stimuli to produce an equal amount of bi-directional interference. Instead, we observed that only average size judgement was very consistently affected by numerosity, suggesting that the principles governing interference for symbolic number-size tasks do not apply in the same way to non-symbolic quantitative stimuli.

The fact that interference is nevertheless more pronounced during mean size judgments, could mean that irrespective of the matched objective degree of discriminability, numerosity has a higher intrinsic salience or capacity to grab attention, and is therefore exerting an influence on response selection. Alternatively, interference might arise from the sensory mechanisms responsible for extracting mean size. Several lines of evidence suggest that mean size is a basic, automatically encoded visual dimension (Ariely, 2001; Chong and Treisman, 2005, 2003; Corbett et al., 2012), which is susceptible to adaptation (Corbett et al., 2012), as numerosity (Burr and Ross, 2008; Ross, 2010). Mean size is thought to be perceived holistically (Ariely, 2001; Chong and Treisman, 2003) through some kind of summary statistics extracted from the visual scene, most likely related to texture rather than individual object processing (Im and Halberda, 2013). Nevertheless, the precise implementation of mean size estimation is currently unknown. Of note, however, Dakin et al. (2011) provided an illustration of how a particular combination of spatial filters applied to an image could provide information about mean item size. Whether this or other similar measures could explain the existence of perceptual biases for mean size, and if so in which direction, will be an interesting question for future studies.

Only very few studies in addition to ours so far investigated the discrimination of numerosity in adult dyscalculic subjects and found that the deficit in non-symbolic numerical proficiency persisted into adult age (Cappelletti et al., 2014a; Cappelletti and Price, 2014; De Visscher et al., 2017; Gilaie-Dotan et al., 2014; Mejias et al., 2012). Here we found that the weber fraction for numerosity was on average lower in dyscalculics than in controls, however this difference did not reach statistical significance, which could be due to the modest sample size available. It is further possible that in adult subjects the non-symbolic enumeration difficulty is more subtle than in children, and easily detected only with more difficult tasks, such as the estimation task used by Mejias et al (2012) or discrimination tasks with displays of spatially intermixed differently colored dots (Cappelletti et al., 2014a; Cappelletti and Price, 2014; Gilaie-Dotan et al., 2014) or sequential presentation (De Visscher et al., 2017) which might exert more demands on working memory compared to the tasks used here. Nevertheless, even in our experiment, we measured a significantly lower accuracy in dyscalculics with respect to controls for the most difficult numerical ratios and, at this level, congruency effects on accuracy were strongest.

Recent studies investigating dyscalculic children or inter-individual differences in the developing population have concluded that enhanced behavioral interference from covarying quantities during numerosity processing are indicative of an impairment of general executive - inhibitory skills which would fully explain the relationship between the approximate number system and math (Bugden and Ansari, 2016; Fuhs and McNeil, 2013; Gilmore et al., 2013; Szűcs et al., 2013). Nevertheless, at least two studies also reported that mathematical competence was associated with numerical acuity over and above inhibitory skills in normally developing children (Bellon et al., 2016; Keller and Libertus, 2015). Our results are in line with the latter findings, as mathematical performance in our group of subjects was correlated with the precision of numerical judgments, even after controlling for inhibitory control, as measured by the color-word Stroop task.

Furthermore, in our psychophysical testing with two different tasks on an equivalent stimulus set, the dyscalculic group showed stronger interference from the unattended dimension than the control group during numerosity judgement only (and not during size judgement). These results are hard to reconcile with the idea of a general inhibition impairment as the source of the interference during quantity judgement, since such an impairment would have been expected to affect both tasks equally.

We do not deny the existence of potential inhibitory deficits in dyscalculia, nor that inhibitory skills play an important role in arithmetic performance in general. Indeed, arithmetic is a complex skill involving a variety of executive attention processes, as well as working memory, fact retrieval, and procedure application. What we are cautioning against here is the uncritical equation of any enhanced interference during quantity processing with a domain general executive function (inhibition) impairment. The enhanced interference during numerosity judgments observed in our dyscalculic group could reflect a difficulty in inhibiting or filtering out irrelevant information which, however, occurs only during numerosity judgments and therefore needs to be domain specific or a heuristic use of non-numerical features to cope with the difficulty in discriminating numbers. Hints in support of the second hypothesis arise from the observation that the direction of interference during numerical judgments was not always the same across subjects in the dyscalculic group, suggesting the adoption of a ‘cognitive’ strategy to solve a task difficult for them.

Indeed, a likely possibility is that these subjects, due to a more imprecise representation of discrete numbers of items, gave more weight in their decisions to low-level dimensions which are partially correlated with numerosity under everyday situations. For example, overestimating numerosities with *larger* dot sizes could indicate some reliance on the overall amount of stimulus energy / total surface area. For overestimation of numerosity with *smaller* dot sizes, it is much less evident which dimension might be relied on. However, this is a common pattern of the interference observed in multiple prior studies in normal subjects, at least for numerosities larger than those used in our study (Gebuis and Reynvoet, 2012a, 2012b; Ginsburg and Nicholls, 1988; Sophian and Chu, 2008; Tokita and Ishiguchi, 2010). Interestingly, this is also the direction of bias predicted by a model based on measures of the relative amount of energy in high and low spatial frequencies of the image (Dakin et al., 2011; Tibber et al., 2012), suggesting that this pattern could be related to the reliance on a texture-like representation of the input.

Furthermore, differences in the direction of the interference (over- as opposed to underestimation) have also been observed previously in normal subjects between different participants within the same study (DeWind et al., 2015, Fig 3). That study used an elegant approach based on a stimulus space which orthogonalized numerosity with respect to two other mathematically derived dimensions (“size in area”, a combination of total surface and individual item area, and “spacing”, an equivalent combination of total field area and sparsity). Their procedure then allowed the authors to determine which of those three main dimensions (or their combinations) best explained subjects’ choices. The intention of our study was somewhat different from theirs: we wanted to evaluate the degree of interference when subjects judge our stimuli on either dimension, rather than numerosity only, as done by Dewind et al (2015). This is why we chose (mean) item size as the dimension orthogonal to numerosity, rather than a dimension such as “size in area” which does not correspond to a natural perceptual dimension that subjects are used to judge. However, this different choice also implies that our design is less suitable for analyses similar to those performed by Dewind and colleagues.

In line with the idea that behavioral interference increases when judgment of the attended dimension becomes more difficult for a subject, we observed a strong correlation between biases in the subjects’ responses and their overall precision during average size discrimination. In other words, the subjects that were less accurate in judging average size were also those showing stronger numerical interference. The same relation appeared for numerical judgments, in which case it coincided with a group effect, with dyscalculics showing lower JNDs and a stronger bias than controls. Crucially, only size interference in numerical judgments correlated with mathematical abilities (even when controlling for the factor of group), supporting a critical link between mathematical performance and numerosity representation specifically, rather than either a general tendency for bias in the presence of incongruency, or the representation of any quantitative dimension.

The fact that here we did not observe size perception to be related to mathematical abilities also fits with other results demonstrating that dyscalculics are not impaired in the discrimination of line length (Cappelletti et al., 2014a; De Visscher et al., 2017) or cumulative area (Iuculano et al., 2008), that education selectively sharpens acuity for numerosity but not single object size (Piazza et al., 2013), and that in the normal population mathematical ability correlates with number, but not with size discrimination thresholds (Anobile et al., 2017), though see (Lourenco et al., 2016) for a significant finding regarding cumulative area. Previous work has also found numerosity but not density sensitivity to be related to the normal development of mathematical abilities in children (Anobile et al., 2016a). Given that average size as density perception is thought to rely on texture processing mechanisms rather than processing of individual items (Im and Halberda, 2013), our findings suggest that texture processing abilities may be preserved in dyscalculics, a possibility that should be further addressed in future studies.

To conclude, using a stimulus set which tested for the amount of mutual interference between numerosity and another quantitative dimension (average item size), with task relevant dimensions matched for discriminability, we found that numerosity could be perceived by normal subjects without significant interference from the irrelevant size dimension. Perhaps more counter-intuitively, mean size was more subject to interference than numerosity in this situation. These results further underline the complex nature of behavioral interference effects between different quantities. More detailed quantitative modelling of how representations of different quantitative dimensions could be derived from the retinal image, or how some dimensions may act as priors modulating perceptual decisions on other dimensions, may help in the future to more fully account for these phenomena. The pattern of interference observed in dyscalculics during the task used here suggest that, in adults at least, enhanced interference during numerosity processing is not the result of a general impairment in executive functions and, more precisely, general inhibitory skills. We propose that these results may reflect the heuristic use of associated stimulus dimensions for task purposes in the presence of a less precise representation of discrete numbers of items, in agreement with the ‘number sense deficit’ theory of dyscalculia. An important goal for future studies will be to understand how neuronal representations of different quantitative dimensions are affected in the dyscalculic brain and how this explain the present behavioral findings.

